# The gut microbiome of the Passalid beetle has high cellulolytic potential and constitutes an unrecognized system for production of greenhouse gasses in neotropical forests

**DOI:** 10.1101/2025.05.12.653568

**Authors:** Gabriel Vargas, Adrián A. Pinto-Tomás, Catalina Murillo-Cruz, Miriam Hernandez, Diego Dierick, Carlos Hernandez, Silvia Soto, Ibrahim Zuniga-Chavez, Tijana Glavina del Rio, Susannah G. Tringe, Jon Clardy, David Sherman, Giselle Tamayo-Castillo

**Author notes:** Funding: NIH project: VI: 809-B4278 y 810-B7-A44.

## Abstract

Herbivores, and their associated microbiomes, play a major role in the global carbon cycle. Recalcitrant cellulose molecules are broken down by microorganisms that colonize the herbivore gut, releasing greenhouse gasses in the process. Passalid beetles are tropical herbivores that feed only on decaying wood and present subsocial behavior that may lead to the acquisition and sharing of microbial symbionts for efficient biomass and energy production. We collected five groups of *Veturius* sp. Passalid beetles from different logs in the Costa Rican rainforest and analyzed the microbial communities of larval and adult guts, as well as the substrate material in which they resided (partially chewed wood material mixed with feces that covers their tunnels). Adults, larvae, and substrate harbor different microbial communities, with substrate showing the highest diversity and richness, and larvae gut comprising a high abundance of methanogenic archaea. Reconstructed metagenome assembled genomes (MAGs) revealed that larvae and adults are enriched in genomes encoding a myriad of glycosyl hydrolases. Methanogenic taxa were enriched within the larval MAGs and bins, suggesting that larval microbiota participate in the final steps of cellulose decomposition in the system. Finally, we assessed methane production rates by Passalid beetles and compared them with termites. Our results show that Passalid larvae and adults produce methane at rates comparable to termites. Passalid and other beetle larvae are potentially underappreciated contributors to the carbon cycle and the biotic production of the greenhouse gas methane.

## Background

Herbivores have evolved different yet efficient strategies such as the evolution of rumen and resistance against plant defenses to decompose plant material and play an important role in the global carbon cycle. The final steps of the remineralization of organic matter produce the greenhouse gasses CO_2_ and CH_4_^1,2^. Methane has 25 times the warming potential of CO_2_ by mass and in nature is produced by anaerobic archaea^3,4^. It has been estimated that archaea produce one gigaton of methane each year in anoxic environments representing about 2% of the CO_2_ fixed annually via photosynthesis^5^.

Invertebrate herbivores including termites and beetles are abundant and widespread in tropical forests and efficient agents of degradation of recalcitrant plant material thus significantly contributing to the carbon cycle^6–8^. The ability of insects to break down cellulose to produce readily accessible carbon molecules depends on the presence of a gut microbial community that harbors functions to hydrolyze cellulose as well as its many degradation products ^6,9–11^. Insects acquire their microbiota mainly from the materials on which they feed; and many of these microorganisms have adapted to the digestive tract environment. Over time, mutually beneficial relationships have developed that enable the hosts to utilize nutrients that would otherwise be unavailable^7,12,13^. Much research involving invertebrate biomass degradation has focused on social insects. Higher termites *Nasutitermes* and *Amitermes* host in their paunch (P3 segment) Spirochaetota and Fibrobacterota that encode hydrolytic enzymes for the degradation of lignocellulose as well as nitrogen fixing bacteria including several Pseudomonadota^6,8,14–16^. The adults of the wood feeding beetle *Odontotaenius disjunctus* harbor beneficial microbes in different anatomical compartments of their gut to facilitate cellulose breakdown^17^. In one study of the microbial community of the wood-boring beetle *Anoplophora glabripennis,* investigators demonstrated the acquisition of specific bacterial groups from the environment, as well as through vertical transmission. These bacteria were involved in mediating nutrient acquisition, nitrogen metabolism and other important processes ^11,16^. However, little research has been done on the microbial communities associated with beetle larvae.

The Passalidae Family includes about 650 species of beetles^18^, and are found from the southern parts of the United States, down to northern Argentina and Uruguay^19,20^. Passalidae consist on approximately 1000 species, and its richness and diversity is the highest in the neotropical forest^21^. They colonize decomposing logs in which they create tunnels for habitation, finding protection from predators and an abundant source of food. This family of beetles is particularly interesting from an ecological point of view as they exhibit eusocial behavior. Traits include the presence of family groups inside logs with overlapping generations and the construction of specific tunnels for laying eggs ^22^. The tunnels made by the beetle are covered in chewed wood material and feces from the adults and larvae, which represents a potential vehicle for the horizontal transmission of beneficial microbes among adults and larvae. In this study we used amplicon sequencing to explore the microbial communities in substrate and the gut of adults and larvae of Passalid beetles in Costa Rica. In addition, we used shotgun metagenomics aiming to reconstruct the genomes of the most important members of the bacterial community to determine their roles in carbon transformations, and the ecology and physiology of the beetle. Finally, we confirmed the production capacity of methane in the Passalid beetle microbiome.

## Methods

### Specimen collection

We collected the beetle specimens for metagenomic DNA extraction in the Braulio Carrillo National Park in Costa Rica: Quebrada González sector, Heredia Province (10° 9′ 36″ N, 83° 58′ 28″ W) in October of 2010. We sampled the interior of decomposing logs and collected larvae and adults from Passalid beetles in the genus *Veturius* sp. This taxonomic identification was confirmed by experts at the Costa Rican National Biodiversity Institute (INBio). All insects were transported to the laboratory at ambient temperature inside vented polyethylene containers and surrounded by the woody material from which they were collected. Additionally, we collected 1g - 5g of the substrate in 50mL Falcon tubes for metagenomic DNA extraction. Once in the lab, all insects were confirmed to be alive and were dissected under sterile conditions, using scissors and forceps to extract the digestive tract for the extraction of genomic DNA. The specimens for methane production assays were collected at La Selva Biological Station in Sarapiquí, Heredia Province, northeastern Costa Rica (10°26′ N, 83°59′ W), during two collection campaigns conducted in August and November 2019. We collected a total of 40 specimens divided in two groups. For Passalid beetles, we collected 10 larvae, 10 adults and 10 substrate samples (∼1g) from several family groups. For comparison, we collected 10 pooled samples of *Nasutitermes* sp. workers (∼0.2g, about 50 individuals) from several colonies. All the individuals were alive upon arrival to the lab.

The collection permits for this work were granted by the Costa Rican Commission for Biodiversity management (CONAGEBIO), the University of Costa Rica Biodiversity Commission and La Selva Biological Station.

### DNA preparation and sequencing

DNA extractions were performed on 69 samples using three commercial DNA extraction kits: FastDNA Spin Kit for Soil (MP), PowerSoil kit (MoBio) and Epicentre MasterPure (MP) with minor modifications to the manufacturer’s protocols. These modifications consisted of different incubation times and the use of previously heated elution buffers to increase extraction yields. Adult and larval gut samples were processed with and without a previous sonication step to test whether sonication facilitates the extraction of DNA from microorganisms that are strongly attached to the gut tissue. None of the substrate samples were sonicated because these lack animal tissues. DNA samples were shipped on dry ice to the DOE Joint Genome Institute (JGI) in Walnut Creek, California for 16S rRNA amplicon sequencing (454 Titanium FLX) and shotgun metagenomic sequencing (Illumina HiSeq 2000). Supplementary Table 1 has a summary of samples, treatments, and DNA extraction methods utilized in this work. The raw amplicons data is available under Bioproject PRJNA1228160, accession SRR32597969.

We first analyzed the diversity of 16S rRNA amplicons (Supplementary Table 2) aiming to determine any differences inherent to the DNA extraction methods and sonication treatment, or any other sampling procedure such as the presence of gut tissue (epithelium). After performing this preliminary analysis of the microbial community composition for all 69 samples, comparing their diversity and richness based on 16S rRNA amplicons, we selected seven samples whose DNA was extracted with the MoBio extraction kit (which provided consistent results in terms of yields) for metagenomic sequencing (Illumina shotgun). These samples included the gut microbiomes of five individual larvae (larvae from families 1-5), one pooled sample from two adult guts (family groups 3 and 4) and one pooled sample from the substrate associated with these insects (from family groups 3 and 4 as well). For this work, we chose the samples extracted with MP protocol for amplicon analysis, due to their higher number of ASVs, and the MoBio extracted samples for metagenomics, due to their higher DNA yield and consistent use of this kit in metagenomic studies.

### Microbiome analysis

The 16S rRNA amplicons were quality filtered and processed using QIIME^23^ and QIIME2^24^ following the recommended workflow. Briefly, we extracted the fasta and quality files from the raw sff files using the script process_sff.py in QIIME. Reads were processed, and quality filtered using split_libraries_fastq.py, imported into Qiime2 and demultiplexed and dereplicated using Cutadapt^25^ and VSEARCH^26^, respectively. Chimeras were detected with UCHIME^27^ and removed from the dataset. The final quality control and the construction of the Feature table was performed with Deblur.

An alignment of the sequences was obtained using MAFFT^28^ and the phylogenetic tree was created using Fasttree^29^ as implemented in QIIME2. The sequences were taxonomically classified using QIIME2 classifier gg-13-8-99-nb-classifier.qza. Finally, the feature table, metadata, taxonomy files and phylogenetic tree were imported into the R package Phyloseq^30^ for further analysis and plotting. First, the sequences classified as Eukaryotic, mitochondria and chloroplast were removed from the data set. Then, the samples were randomly rarefied to the smallest sample size = 6412 sequences (except sample 5MSA which only had 34 sequences and was removed from the analysis). Alpha and beta diversity, as well as barplots and statistics, were calculated and plotted using the Phyloseq, Vegan, ggplot2, Microbiome, and Phylosmith R packages.

The metagenomic Illumina reads were quality filtered using Cutadapt^25^, and the remaining reads were co-assembled using Megahit^31^ (kmer sizes from 31 – 91 nucleotides). The metagenomic analysis was performed following the Anvio v6.2^32^ pipeline. Briefly, the filtered reads were mapped back to the co-assembly with Bowtie2^33^ and Samtools^34^ in order to calculate the coverage of the contigs in the metagenomes. The bam files were indexed in Anvio v6.2 and used to create a Profile database of the co-assembly. A contigs database was generated in Anvio as well. The metagenomic contigs were then binned using Metabat2^35^, and imported into the Anvio databases. Finally, we employed Anvio’s interactive tools to explore and visualize metagenomes, to manually refine metagenomic bins, and to evaluate reconstructed metagenome assembled genomes (MAGs). The quality of the metagenomic bins was assessed using checkM^36^. We defined MAGs as follows: bins larger than 1Mb with >70% completion and <7% redundancy. Open reading frames (ORFs) were called for the MAGs using Prodigal^37^. Taxonomical and functional annotations were performed against the databases Clusters of Orthologous Groups^38^ (COG) and Carbohydrate-Active Enzymes^39^ (CAZY) with Diamond^40^ using the sensitive setting. The taxonomic classification of MAGs was performed using GTDB-tk^41^. The phylogenomic trees were constructed following Anviós v6.2 protocols available online and edited in Interactive Tree of Life (iTOL).

The metagenomes were also submitted to the Integrated Microbial Genomes and Metagenomes (IMG/M) and analyzed using their pipeline. They were assembled using SOAPdenovo, Newbler and Minimus2, and annotated against functional databases. These data is available under JGI Project IDs: 404336, 404655-303660.

### Methane production measurements

Whole individual insects were used for methane production assays. Specimens were weighed to determine their fresh weight and introduced in a suitably sized vials (20-60ml) sealed with a rubber septum for 120 minutes incubation. Blank samples were included as controls and to determine CH_4_ concentration at the start of the incubation. After the incubation period, gas was extracted from the sample vials and analyzed for CH_4_ on a 7890B gas chromatograph (Agilent Technologies, Santa Clara, California, USA) equipped with FID and a HP-PLOT/Q column (30m, 0.53mm, 40um). The measured CH_4_ concentrations were then used to estimate CH_4_ production rates (fCH_4_, nmol*g-1*h-1) based on sample mass, incubation time and initial CH_4_ levels. The results were plotted using ggplot2 and ggpubr R packages.

## Results

### Community diversity and composition

We collected five Passalid beetle family groups from individual decaying logs found inside their characteristic tunnels. These tunnels are covered with a substrate (Fig1C). We initially hypothesized that the substrate is an important vehicle for microbial transmission between adults and larvae. Hence, we sequenced the gut contents of Passalid adults and larvae as well as the substrate corresponding to each beetle family collected in this experiment. The guts of the adults and larvae are anatomically different (Figure 1B, 1D respectively), and contain different gut contents. The adult gut is compartmentalized as it has been described in other adult insects, and the contents consisted of a mixture of wet and dark wood materials, suggesting that these were subject to chemical and enzymatic transformations. On the other hand, the larval gut consists of a straight tube with no evidence of compartmentalization based on stereoscopic inspection. The contents of the larval gut also differed from the adults as these were dryer and less mixed, like the substrate material, in which it was still possible to recognize wood fibers.

**Figure 1.**
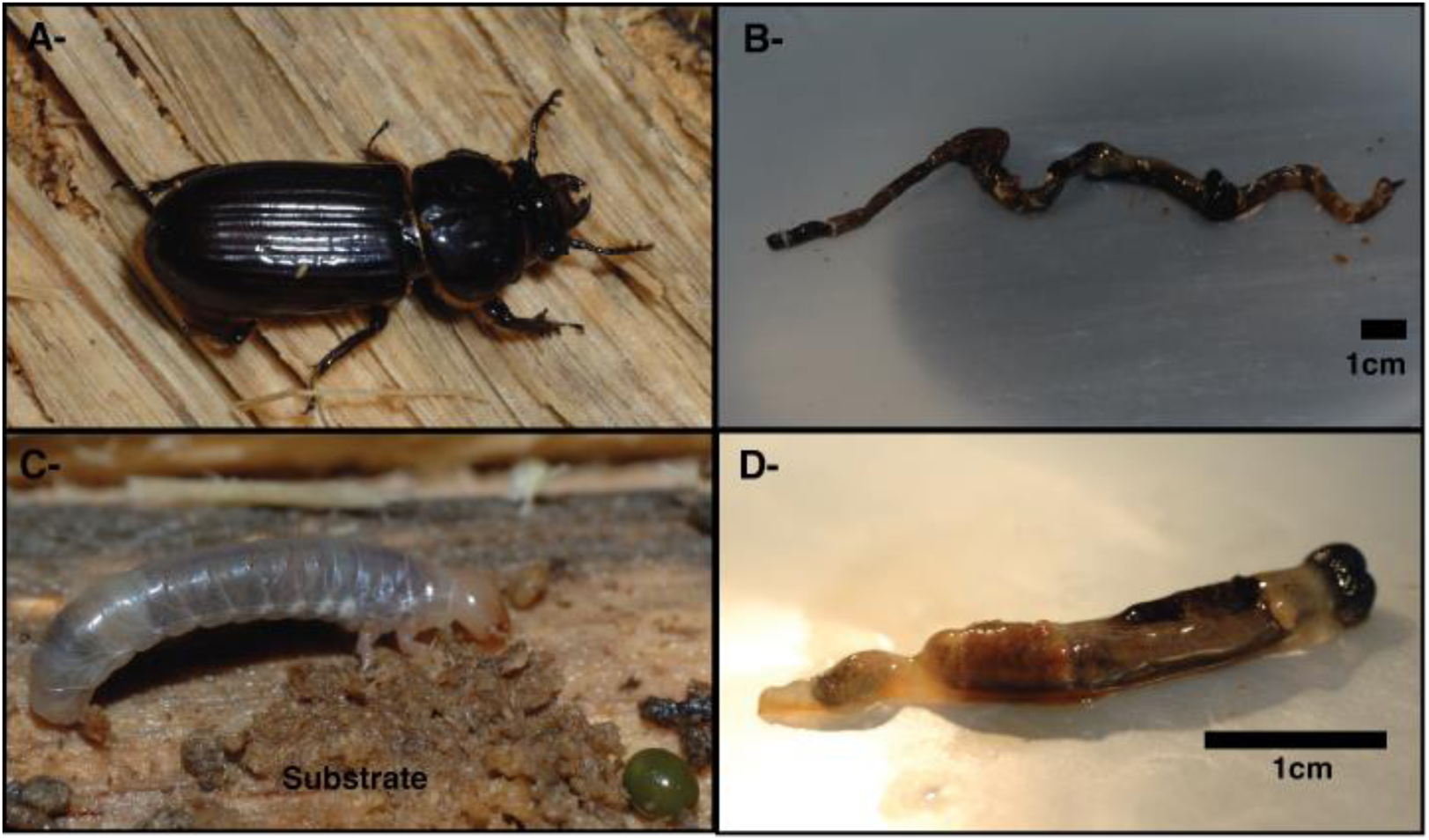
Passalid beetle eusocial system. A-Adult *Veturius* sp. beetle inside an uncovered decaying log and surrounded by the substrate. B-Dissected gut of the adult *Veturius* sp. C-Larvae of *Veturius* sp. inside an uncovered decaying log and surrounded by the substrate (GM) and a green egg. D-Dissected gut of the *Veturius* sp. larvae.

The diversity of the samples was evaluated using alpha diversity metrics for richness (Observed and Chao1), Shannon and Inverse Simpson (Supplementary Table 2). There were no significant effects of the DNA extraction method or the sonication treatment. Based on these results, we continued the analysis using a subset of 15 DNA samples that represented the five Passalid beetle families collected from the five different logs. The substrate samples showed the highest richness and diversity compared to the adult and larval gut (Fig. 2AB), and sample type (adult, larvae and substrate) strongly determines the composition of the microbial communities (Figure 2CD; ANOSIM= 0.784, p value= 0.001). This grouping pattern is stronger when taking abundance of taxa into account using weighted Unifrac in which the two first components explain 83.6% of the variation (Fig. 2D). When abundance is not considered (unweighted Unifrac), the grouping pattern remains consistent, but the first two components explain only 31% of the total variation (Fig. 2C). By contrast, other variables such as the family group (ANOSIM= -0.109, p value= 0.789), are not significant contributors to the variation. In this analysis the variable family group comprises the effect of the variables: small-scale geographical location, type of wood in which the beetles were collected, and other environmental factors specific to each sample.

**Figure 2.**
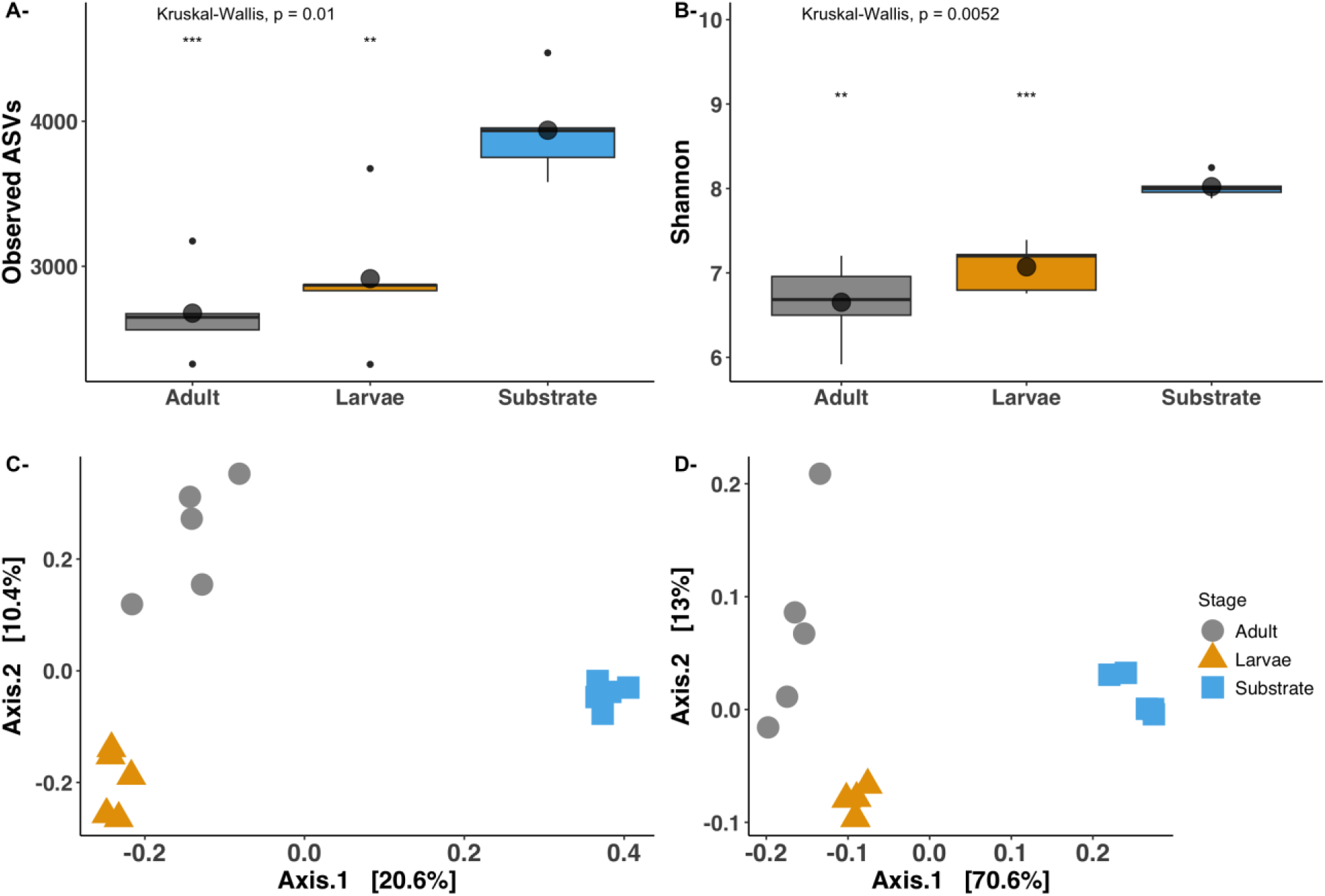
Alpha and beta diversity of the Passalid beetle microbiome. A- and B-Alpha diversity metrics: Observed ASVs and Shannon metrics respectively. Kruskal Wallis was used for the overall comparison of the means (represented with circles). Adult and larvae mean were compared to the substrate mean using Wilcoxon test. **= p<0.01, ***= p<0.001. C-Unweighted Unifrac PCoA. D-Weighted Unifrac PCoA. Gray= adults. Yellow= larvae. Blue= substrate. The samples were rarified to 6412 reads. The numbers represent the five families. BSU= substrate, BSA= adults and BSL= Larvae.

The microbial community composition is different for all three sample types (Fig. 2). The taxonomic distribution at phylum level shows that the substrate samples are largely dominated by members of Pseudomonadota followed by Bacteroidota, Acidobacteriota and Verrucomicrobiota. In the case of the larval gut, these microbial communities are dominated by the archaeal phylum Methanobacteriota, followed by Bacillota and Bacteroidota. Finally, in the adult gut microbial communities the predominant phyla were Bacillota and Mycoplasmatota, followed by the Bacteroidota and Pseudomonadota (Fig. 3A).

**Figure 3.**
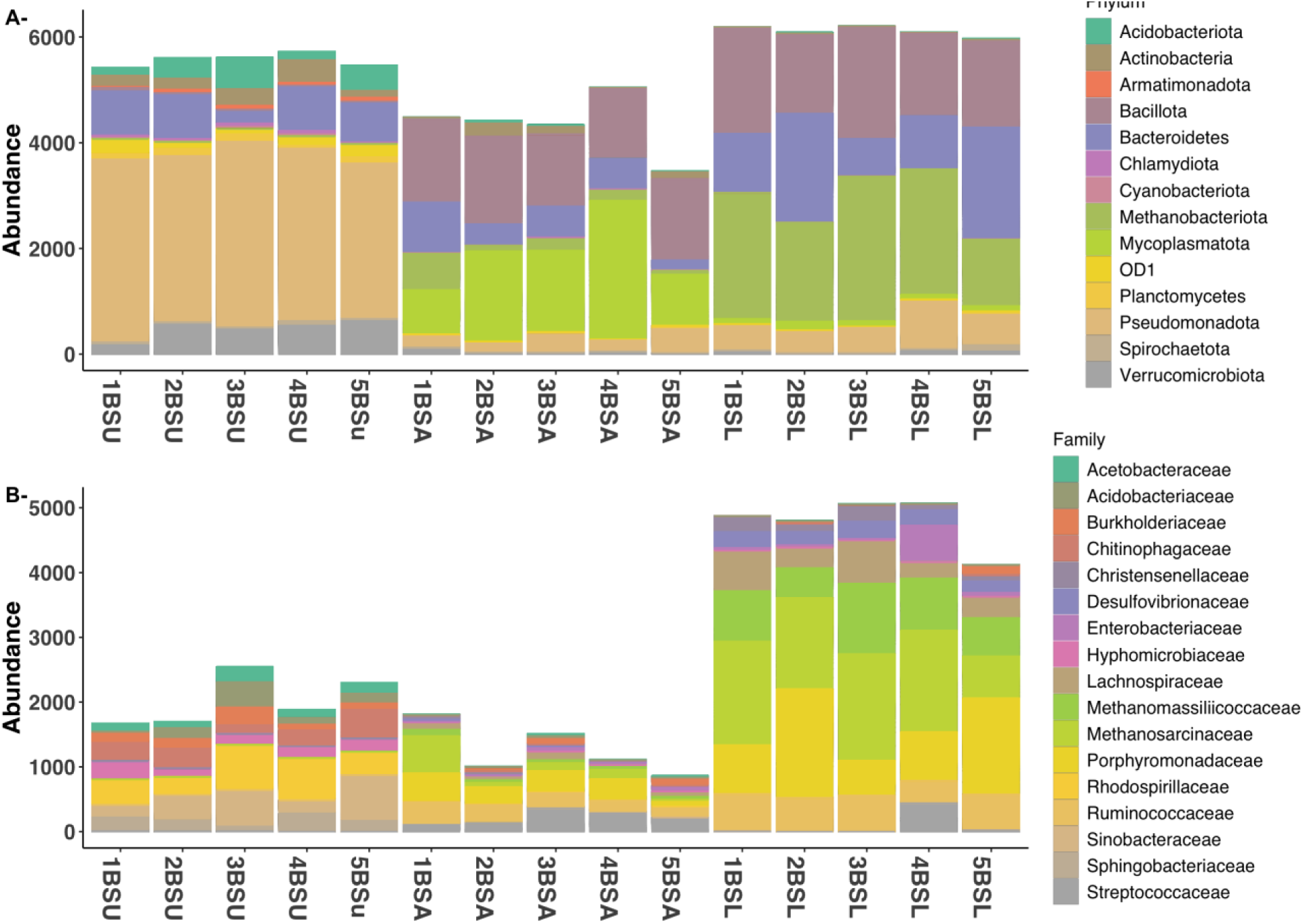
Phylogenetic distribution of the Passalid beetle microbiome. A) Phylum level distribution of the most abundant phyla (>1% relative abundance representing 82.4% of all ASV). B) Family level distribution of the most abundant families (>1% relative abundance representing 68.1% of all ASV). The samples were rarified to 6412 reads. 13.4% of the ASV were assigned only classified to Bacteria. The numbers represent the five beetle families. BSU= substrate, BSA= adults and BSL= Larvae.

Bacillota families Lachnospiraceae, Streptococcaceae, Clostridriaceae and Ruminococcaeae are within the 14 most abundant families in the entire dataset (Fig. 3B). The archaeal phylum Methanobacteriota showed the highest abundance in the gut of Passalid larvae, specifically the methanogenic families Methanosarcinaceae and Methanomassiliicoccaceae, which are the two most abundant families in larval samples. The phylum Mycoplasmatota was abundant in the adult gut compared to the larvae or substrate samples. Among the Pseudomonadota, several families of the class Alpha Proteobacteria showed high abundance in the substrate including Rhodospirillaceae, Hyphomicrobiaceae and Enterobacteriaceae. The phyla Bacillota, Mycoplasmatota and Methanobacteriota are the most important contributors to the differences observed in the composition of the gut microbial communities in comparison to the substrate. The most abundant microbial families represented between 80 – 95% of their entire community. This is much higher than what it was observed in the adult and substrate microbial communities (Fig 2B).

### Metagenomic analysis

We focused the shotgun sequencing efforts on the larvae gut due to its high abundance of archaea and other bacterial groups of interest. The adult and substrate samples were sequenced as a pool of the samples from two families of beetles. The co-assembly of the metagenomes produced 2.7 Gb worth of contigs with the longest one being 413kb and N50 of 23.8kb (contigs smaller than 1kb were excluded from the analysis). Remaining contigs were reconstructed/assembled into 766 bins, 391 of which were larger than 1Mb. The average bin size was 1.3Mb and the average number of ORFs obtained per bin was 1290. The average %GC of the bin collection was 46.52%, presenting a slight bias towards low GC-content microorganisms. Among all bins, 92 fulfilled our selection criteria and were therefore considered as metagenome assembled genomes (MAGs). Figure 4 shows the distribution of MAGs in a phylogenomic tree. Most MAGs and bins were mapped back to the larvae metagenomes (Sup. Fig. 1). Of the 92 MAGs, 59 were classified as Bacillota, followed by Bacteroidetes (8), Verrucomicrobia (7) and Pseudomonadota (6) (Sup. Table 3). In total, we obtained MAGs from 14 different phyla including 2 archaeal phyla. One of the archaeal MAGs belonged to the class Bathyarchaeia, while the other was classified in the genus *Methanoplasma*.

**Figure 4.**
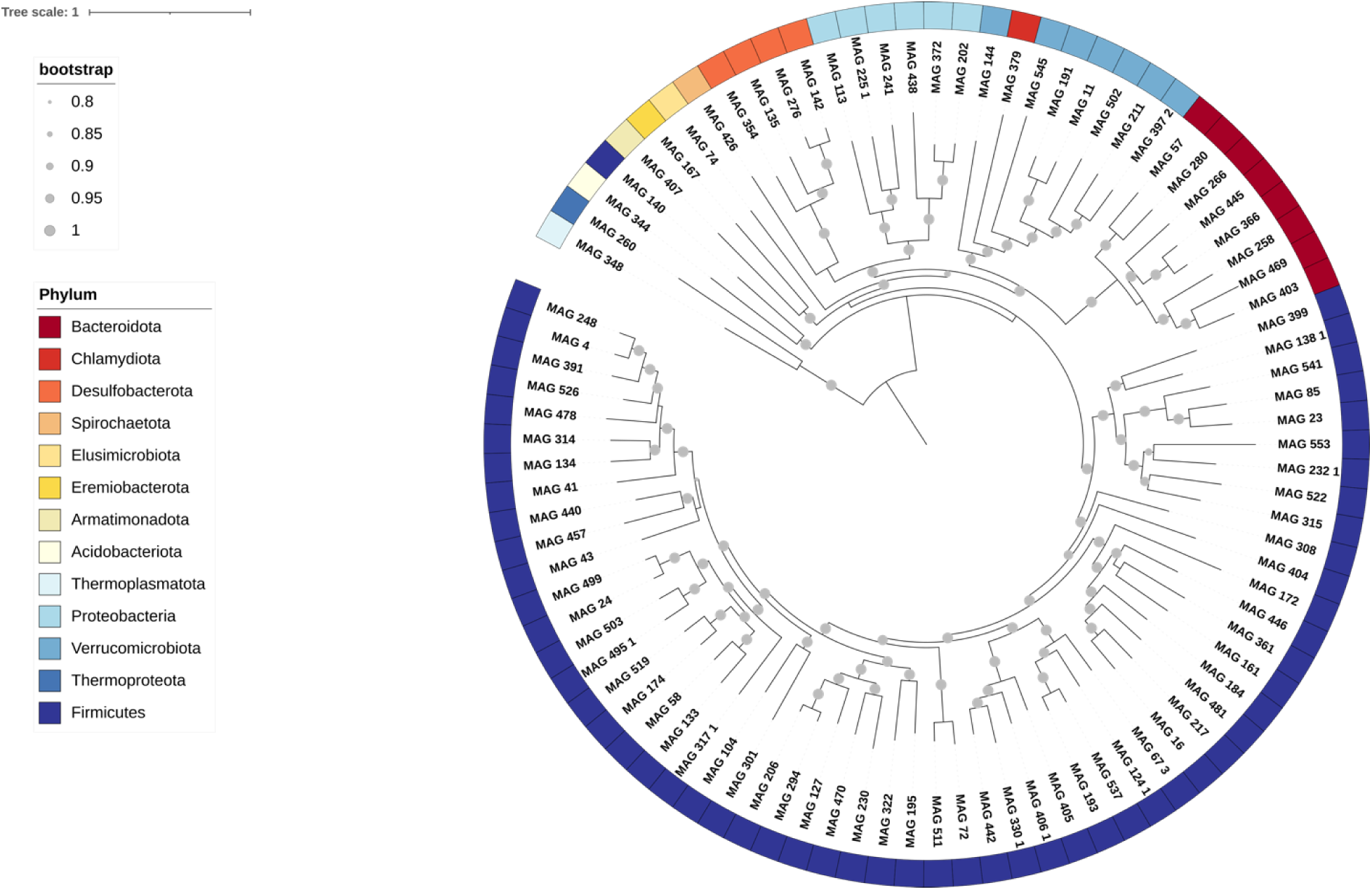
Diversity of MAGs reconstructed from Passalid beetle metagenomes. Phylogenomic tree constructed using 71 concatenated single copy core genes. Colors represent phyla. Gray circles represent the bootstrap values, only values >0.8 are shown.

### Cellulose degradation in MAGs

Given the importance of efficient consumption of plant material for Passalid beetles, we analyzed the cellulose degradation capabilities of the 92 reconstructed MAGs. We annotated the predicted ORFs in each MAG against the Carbohydrate-Active enzymes database (CAZy). We found a wide variety of hydrolases and other accessory functions involved in cellulose degradation and other carbon transformations. Figure 5 shows the distribution of glycosyl hydrolases (GH) clans, glycosyl transferases (GT), carbohydrate binding modules (CBM) and carbohydrate esterases (CE) in each MAG. The GH-A clan presented the highest abundance among the glycosyl hydrolases, these include several families of β-glucosidases and β-galactosidases, as well as other glycosyl hydrolases such as GH9 and GH28 families of cellulases. Clans GH-B and GH-H were also highly abundant in the MAGs. The former includes GH7 which cleaves β-1,4 glycosidic bonds in cellulose and has xylanase activity, and GH16 which includes a large variety of hydrolases that act on plant polysaccharides. The latter comprises GH13 which shows activity on polysaccharides such as amylose and amylopectin and the transglycosylases GH70 and GH77. The clan GH-K showed intermediate abundances in the MAGs and contains the family GH18 of chitinases.

**Figure 5.**
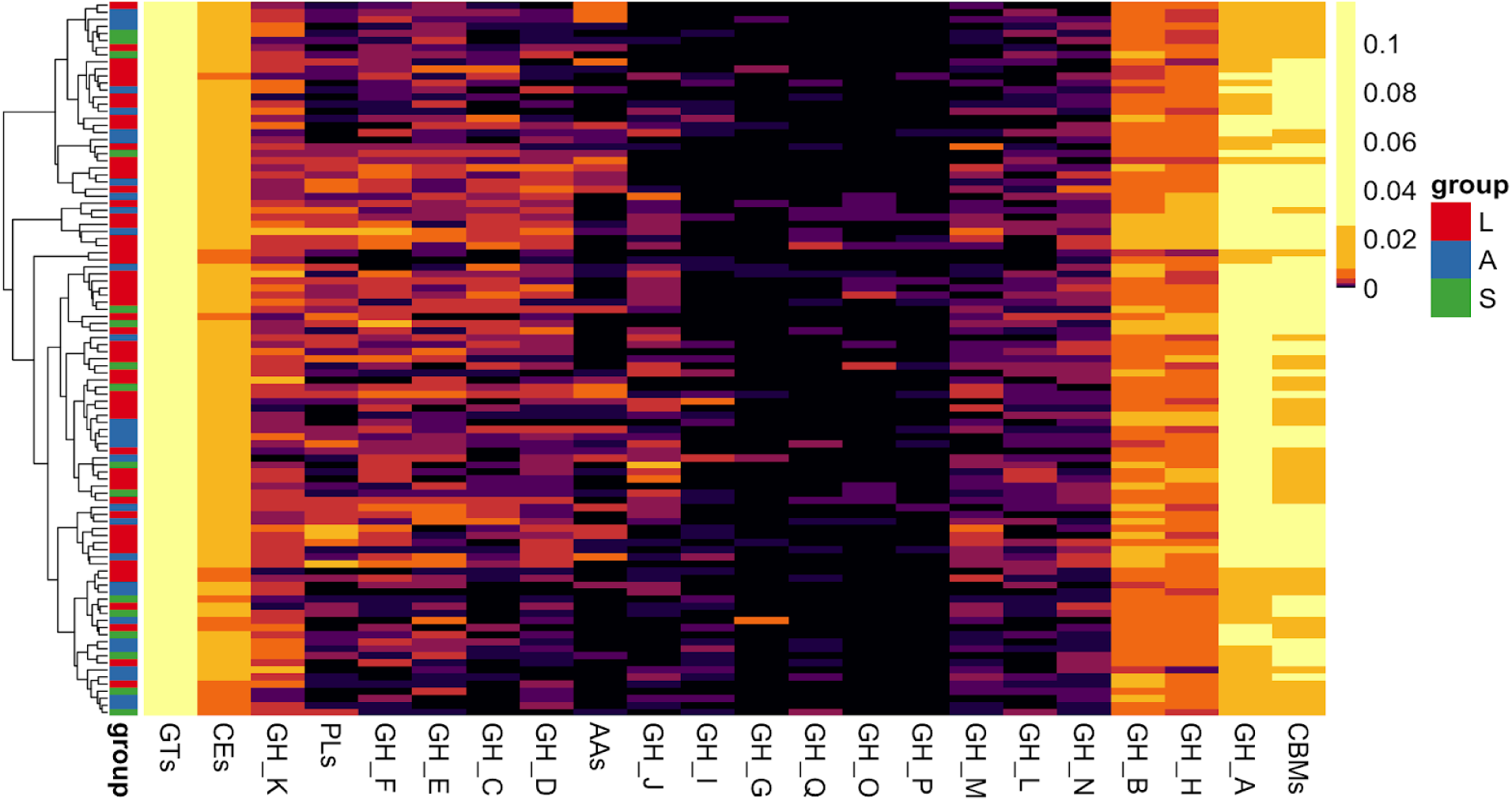
Heatmap of the functions related to cellulose breakdown on reconstructed MAGs from the *Veturius* sp. gut and substrate metagenomes. GH= glycosyl hydrolase, GT= glycosyl transferase, CE= carbohydrate esterase, AAs= auxiliary activities, PL= polysaccharide lyase, CBMs= carbohydrate binding modules. The origin of each MAG is showed in groups: Red (L)= Larvae, Blue (A)= Adult, and Green (S)= Substrate. Data was normalized to MAG gene count. Clustering is based on CAZymes abundance data.

All MAGs also showed significant presence of GTs, CEs and CBMs. GTs are expected to be abundant as they take part in the biosynthesis of oligo and polysaccharides by forming glycosidic bonds; these molecules are metabolically important for biomass and energy production. CEs were consistently abundant in all MAGs, especially in Bacillota. CBMs were also abundant across all MAGs, however, they showed higher abundances in terms of genes and sequences in the larval MAGs. Finally, some auxiliary activities (AAs) families are known to have ligninolytic activity or to act in conjunction with GHs to break down lignin. AAs were significantly less abundant in our MAGs suggesting that lignin degradation is being performed by other microorganisms that we failed to assemble and bin, or that some of the GHs in our MAGs have this activity.

### Methanogenesis in the Passalid beetle system

The phylogenetic analysis based on 16S rRNA sequences showed a large abundance of archaea, especially in the larval gut samples. Hence, we analyzed our metagenomic data to identify archaeal taxa and reconstructed two MAGs. A phylogenomic analysis of MAG260 shows it clustered within other Bathyarchaeia (phylum Thermoproteota) MAGs reconstructed from termite metagenomes (Sup. Fig. 2). These results suggest the existence of a clade of archaea in this class that is associated with herbivorous insects.

Guided by the discovery of the archaea present in the guts of Passalid adults and larvae, we determined the beetle’s ability to produce methane and compared it with other systems known for their methane production. We found that both adults and larvae produce methane at similar rates to what we measured for the higher termite *Nasutitermes* sp. Interestingly, there was no production of methane in the substrate, probably due to the low abundance of archaea in this sample type and its aerobic atmosphere.

## Discussion

In this study we characterized the taxonomic and functional diversity of the microbiome of Passalid beetles from Braulio Carrillo National Park, Costa Rica. The microbial communities in the gut content of the adult and larvae, as well as the associated substrate, are phylogenetically diverse and have distinct community compositions. These are enriched in microorganisms with the genetic potential to play important roles for the community itself and the host, such as cellulose degradation and methanogenesis. Our findings suggest that the gut microbiota play a role in the ecology of these beetles. We hypothesize that they cooperatively degrade cellulose, and that larval and adult guts host methanogenic archaea that complete the catabolic metabolism of carbon to generate methane. This characteristic might be a major driver for the subsocial behavior of these beetles.

One of the most interesting results we obtained was the striking difference between the adult, the larvae and the substrate microbial communities. We found that sample type was the most important driver of the variation of the microbial community composition, overpowering the contribution of the beetle family group and other variables such as geographical location, the tree species in which beetles burrow and environmental factors. The beta diversity clustering observed was supported by the differences in the phylogenetic groups that dominated each sample type. The Passalid substrate samples were largely dominated by Pseudomonadota, the adults by Bacillota and Mycoplasmatota, and the larvae by Methanobacteriota and Bacillota. This is opposite to what as been observed in other beetles. In the wood feeding beetle *Anoplophora glabripennis*, the authors found similar microbial communities between different life stages of the beetle and the wood material in their galleries, and concluded that the microbes in the larval gut were either vertically transmitted or environmentally acquired from the bark they feed on^11^. Two other metagenomic study of the Mountain pine beetle *Dendroctonus ponderosae* and the bark beetle *Dendroctonus valens* LeConte also showed similar microbial communities in the larvae and adults, again dominated by Pseudomonadota^50^. Finally, a study of the communities associated with another Curculionidae beetle *Hylobius abietis,* found that these were very similar in individuals across Europe and dominated by Pseudomonadota followed by Bacillota^51^. In contrast, our results determined that the communities in the Passalid beetles and substrate are different, only sharing a small percentage of the ASVs. Moreover, there are high order differences in the dominant taxa even at the phylum level.

The Passalid adult and larvae guts are enriched in microorganisms capable of cellulose degradation to generate methane and carbon dioxide. Their communities are characterized by Clostridiales including members of Clostridiaceae, Ruminococcaceae and Lachnospiraceae, as well as other Bacilli. The larvae are also rich in methanogenic archaea comprising members of several classes of Methanobacteriota. These results are consistent with those found in other cellulolytic microbial communities such as the cow rumen and the termite hindgut. Several studies of mammalian rumen have identified the Bacillota (formerly Firmicutes) as the taxa carrying the largest and most diverse repertoires of carbohydrate active enzymes^9,52–54^. In a study of the hindgut of two different species of termites, the authors found that the taxa carrying the cellulolytic capacity differ between the species; in *Amitermes wheeleri* the most abundant phyla were the Bacillota and Spirochaetes, whereas in *Nasutitermes corniger* the community was dominated by the Spirochaetes and Fibrobacteres^6,8^. Neither Spirochaetes nor Fibrobacteres showed within the most abundant phyla in the Passalid metagenomes. The cow rumen is rich in methanogenic archaea, most of which are classified as *Methanobrevibacter.* ^55^

We reconstructed several hundred metagenomic bins of microorganisms that inhabit the Passalid gut and its substrate. These are distributed in several bacterial and archaeal phyla. From those, 92 satisfied our definition of MAG, and therefore our research was concentrated in those near complete genomes. Most of the MAGs reconstructed from the Passalid metagenomes correspond to Bacillota and contain many glycosyl hydrolases and other genes for carbohydrate active enzymes, supporting a key role for Bacillota in the degradation of cellulose in the Passalid gut. Others have attempted to assemble genomes from cellulolytic metagenomes, specifically from the cow rumen. In one of the first studies, the authors were able to assemble 15 genomes of uncultured bacteria from the phyla Bacillota, Bacteroidetes and Spirochaetes^52^. In a 2018 study Stewart et al. used a combination of HiSeq Illumina sequencing and Hi-C Illumina sequencing^56^ to reconstruct draft genomes of 898 microorganisms assembled from 43 cow rumen metagenomes. Many of their MAGs were classified in the Bacillota and Bacteroidetes phyla, followed by genomes of Actinobacteria, Pseudomonadota and Methanobacteriota^55^. Their collection resembles ours in terms of taxonomic affiliation of the genomes and in their overall functional capacity. Both rumen and Passalidae datasets showed great novelty in the thousands of sequences that encode carbohydrate metabolism (estimation based on the percentage of sequences with good matches to the public databases), however there is a significant difference in terms of the bacteria that encode these enzymes. While in cow rumen MAGs the majority of the glycosyl hydrolases are detected in Bacteroidetes (Prevotellaceae), in the Passalid beetle these are significantly more abundant in the Bacillota, specifically in the Clostridiales. The approximated taxonomy of the archaea found in both collections is also distinct; 25 of the 28 archaeal genomes in the Stewart dataset were known ruminant *Methanobrevibacter*, whereas our MAGs are in other novel phyla including Thermoproteota. This group is of particular interest as it has been shown to be present in the gut of several termite species^42^. The cow rumen and the Passalid beetle MAGs have the potential to substantially improve our knowledge of microbes specialized to degrade cellulose in the herbivorous gut environment. Many of these microorganisms are unculturable in the laboratory under current culturing methods, making these datasets a valuable resource for bioprospecting purposes.

Wood feeding animals rely on microbes to degrade recalcitrant cellulose and lignin molecules into more easily digestible compounds for the host. Therefore, the microbial communities in the gut of Passalid beetles, and other animals, heavily influence their ecology in terms of habitat selection, diet and behavior. The Passalid adult and larvae have distinct cellulose degradation capacities, suggesting that they use different strategies to produce energy and biomass from this polymer. Based on our findings, we propose a model to explain how the enrichment of specific microbial taxa involved in carbon metabolism in the gut of the Passalid beetles contributes to the evolution of the characteristic subsocial behavior of this family of Coleoptera (Fig. 7). In the model we propose that the larvae are more intensively contributing to the cellulose breakdown in the system. The beetle larvae are very voracious and rapidly growing^18,57,58^, thus the larvae have a high energy demand to produce biomass. Additionally, the larval gut occupies most of its body, consisting of a tube that lacks anatomical compartmentalization. Due to these features, and other direct observations of the integrity of the material inside the larvae gut during the extraction, we believe that the contents transit rapidly through the gut, preventing a complete digestion of the plant material before its ejection in the feces. Both larvae and adults could feed on the easily digestible sugars in the substrate produced by the associated microbial community. Then, inside the insect gut, the Bacillota and methanogenic archaea eventually completely transform cellulose to CO_2_ and CH_4_. The adult metagenomes were significantly depleted in cellulases like GH5, GH10 and others involved in the first steps of cellulose breakdown; this is why we believe they benefit from the partially degraded sugars produced by the larvae. Thus, the subsocial behavior of the Passalids could be at least partially explained by the dynamics of their relationship where the larvae assist with digestion for the adult, and the adult provides the larvae with protection and a habitat.

**Figure 6.**
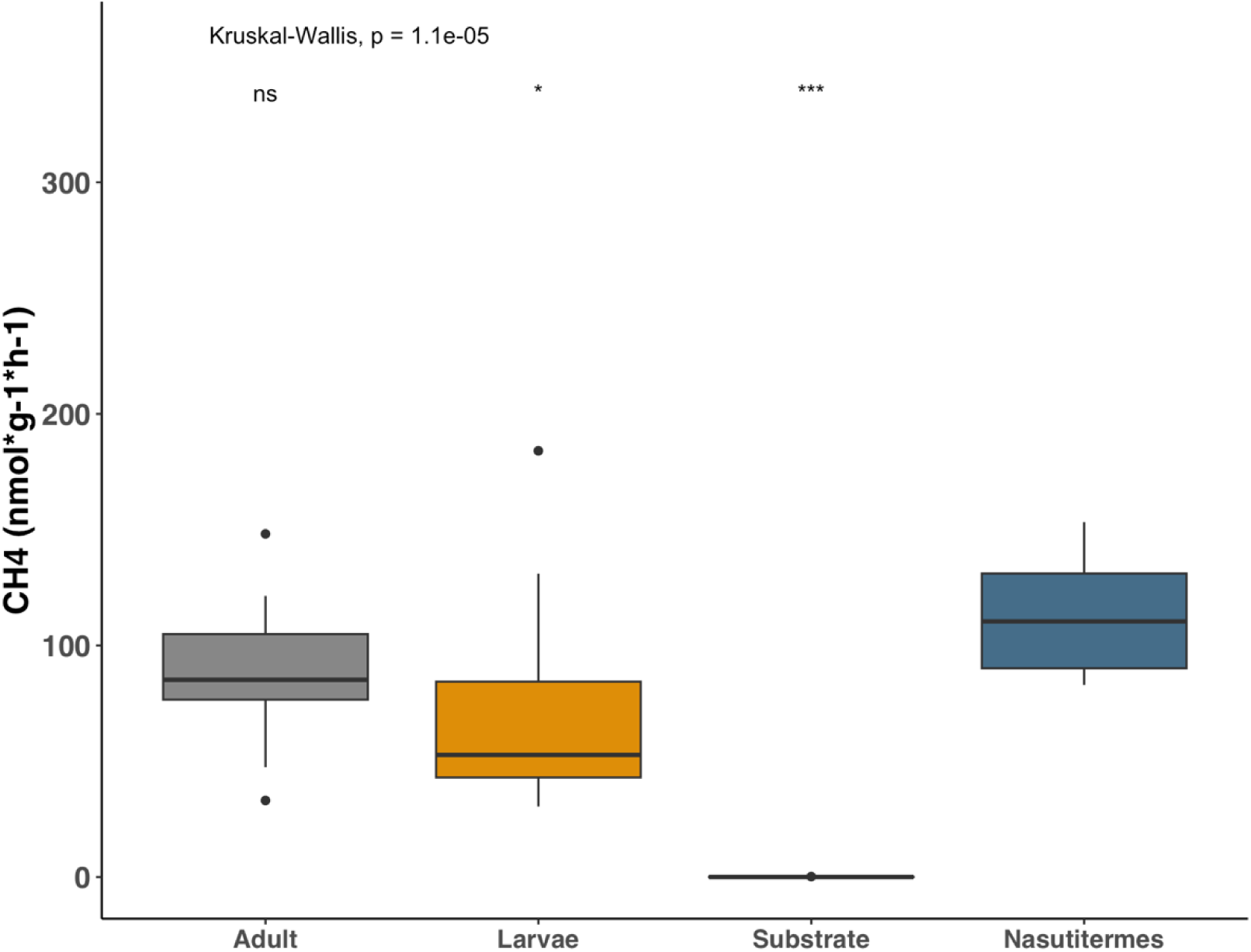
Methane production of whole Passalid beetles and termites. Data was not normally distributed according to a Shapiro test. Overall differences between groups were determined using Kruskal-Wallis test (n=10), Pairwise comparisons performed using the Wilcoxon test and *Nasutitermes* sp. as reference.

**Figure 7.**
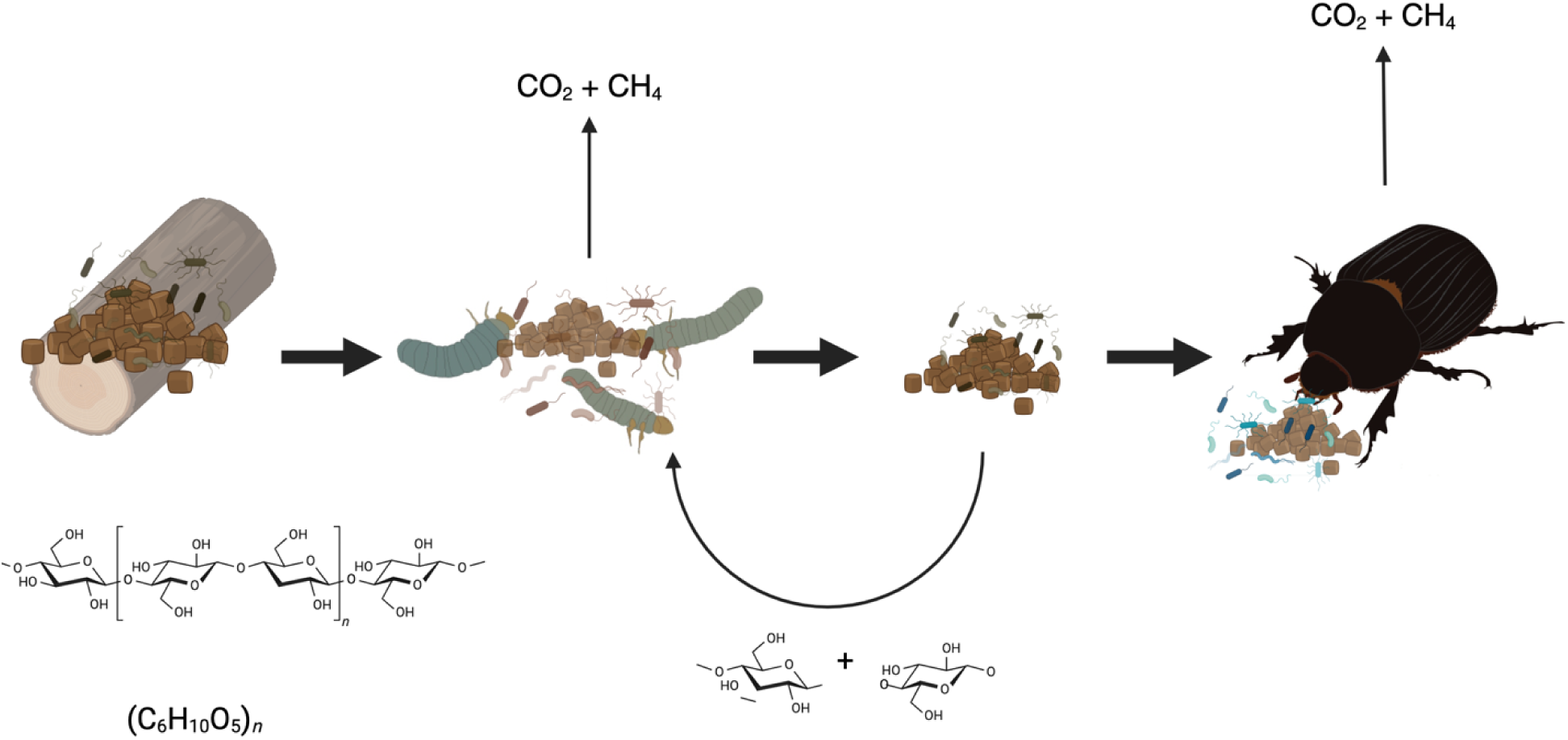
Proposed model of microbe-assisted cellulose degradation in the Passalid beetle *Veturius* sp.

Methane production measurements showed that Passalid beetles, both larvae and adults, produce methane at levels equivalent to what is observed in higher termites. Given the abundance of these beetles in tropical forests, they might be significantly contributing to methane production. It has been estimated that the microbial production of methane is about one billion tons per year^1^. While most methanogenic archaea are thought to reduce CO_2_ with hydrogen to produce methane, our dataset also found evidence for the use of other substrates like formate, methanol, acetate and some new pathways for energy production^2^. The methanogens play an important role in cellulolytic systems dissipating decomposition products, such as methanol and organic acids that can accumulate to toxic levels, releasing them to the atmosphere^44^.

Taken together our results validate the Passalid beetle gut as a phylogenetically and functionally diverse cellulolytic system with characteristics similar to the cow rumen and the termite hindgut. Nevertheless, we also demonstrated the uniqueness of the Passalid microbial community and contributed a dataset of partial genomes that can be further used to better understand the ecology and natural history of these and other beetles in the tropics. Finally, we propose a model to explain the role of the gut microbial community in the ecology and physiology of the Passalid beetle. This model, based on a snapshot of metagenomes and 16S rRNA data from five families of Passalid beetles, will require validation and refinement by future research involving collections from multiple geographical sites, gene expression measurements and controlled laboratory experiments.

## Acknowledgments

The work (proposal: 10.46936/10.25585/60007372) conducted by the U.S. Department of Energy Joint Genome Institute (https://ror.org/04xm1d337), a DOE Office of Science User Facility, is supported by the Office of Science of the U.S. Department of Energy operated under Contract No. DE-AC02-05CH11231.

---

**Supplementary Table 1.** Samples for DNA isolation and 16S rRNA amplicon sequencing.

| **Sample\*** | **Sample Type** | **Stage** | **Treatment** | **Extraction Method** |
| --- | --- | --- | --- | --- |
| 1BA | Gut content | Adult | No sonication | MP |
| 1BL | Gut content | Larvae | No sonication | MP |
| 1BSA | Gut content | Adult | Sonication | MP |
| 1BSL | Gut content | Larvae | Sonication | MP |
| 1BSU | Gut content | Substrate | No sonication | MP |
| 1MA | Gut content | Adult | No sonication | Mobio |
| 1ML | Gut content | Larvae | No sonication | Mobio |
| 1MSA | Gut content | Adult | Sonication | Mobio |
| 1MSL | Gut content | Larvae | Sonication | Mobio |
| 1MSU | Gut content | Substrate | No sonication | Mobio |
| 2BA | Gut content | Adult | No sonication | MP |
| 2BL | Gut content | Larvae | No sonication | MP |
| 2BSA | Gut content | Adult | Sonication | MP |
| 2BSL | Gut content | Larvae | Sonication | MP |
| 2BSU | Gut content | Substrate | No sonication | MP |
| 2MA | Gut content | Adult | No sonication | Mobio |
| 2ML | Gut content | Larvae | No sonication | Mobio |
| 2MSA | Gut content | Adult | Sonication | Mobio |
| 2MSL | Gut content | Larvae | Sonication | Mobio |
| 2MSU | Gut content | Substrate | No sonication | Mobio |
| 3BA | Gut content | Adult | No sonication | MP |
| 3BA | Epithelium | Adult | No sonication | MP |
| 3BepA | Epithelium | Larvae | No sonication | MP |
| 3BL | Gut content | Larvae | No sonication | MP |
| 3BSA | Gut content | Adult | Sonication | MP |
| 3BSL | Gut content | Larvae | Sonication | MP |
| 3BSU | Gut content | Substrate | No sonication | MP |
| 3MA | Gut content | Adult | No sonication | Mobio |
| 3MepA | Epithelium | Adult | No sonication | Mobio |
| 3MepA | Epithelium | Larvae | No sonication | Mobio |
| 3ML | Gut content | Larvae | No sonication | Mobio |
| 3MSA | Gut content | Adult | Sonication | Mobio |
| 3MSL | Gut content | Larvae | Sonication | Mobio |
| 3MSU | Gut content | Substrate | No sonication | Mobio |
| 4BA | Gut content | Adult | No sonication | MP |
| 4BepA | Epithelium | Adult | No sonication | MP |
| 4BepL | Epithelium | Larvae | No sonication | MP |
| 4BL | Gut content | Larvae | No sonication | MP |
| 4BSA | Gut content | Adult | Sonication | MP |
| 4BSL | Gut content | Larvae | Sonication | MP |
| 4BSU | Gut content | Substrate | No sonication | MP |
| 4MA | Gut content | Adult | No sonication | Mobio |
| 4MepA | Epithelium | Adult | No sonication | Mobio |
| 4MepL | Epithelium | Larvae | No sonication | Mobio |
| 4ML | Gut content | Larvae | No sonication | Mobio |
| 4MSA | Gut content | Adult | Sonication | Mobio |
| 4MSL | Gut content | Larvae | Sonication | Mobio |
| 4MSU | Gut content | Substrate | No sonication | Mobio |
| 5BA | Gut content | Adult | No sonication | MP |
| 5BepA | Epithelium | Adult | No sonication | MP |
| 5BepL | Epithelium | Larvae | No sonication | MP |
| 5BL | Gut content | Larvae | No sonication | MP |
| 5BSA | Gut content | Adult | Sonication | MP |
| 5BSL | Gut content | Larvae | Sonication | MP |
| 5BSU | Gut content | Substrate | No sonication | MP |
| 5EA | Gut content | Adult | No sonication | Epicentre |
| 5EepA | Epithelium | Adult | No sonication | Epicentre |
| 5EepL | Epithelium | Larvae | No sonication | Epicentre |
| 5EL | Gut content | Larvae | No sonication | Epicentre |
| 5ESA | Gut content | Adult | Sonication | Epicentre |
| 5ESL | Gut content | Larvae | Sonication | Epicentre |
| 5ESU | Gut content | Substrate | No sonication | Epicentre |
| 5MA | Gut content | Adult | No sonication | Mobio |
| 5MepA | Epithelium | Adult | No sonication | Mobio |
| 5MepL | Epithelium | Larvae | No sonication | Mobio |
| 5ML | Gut content | Larvae | No sonication | Mobio |
| 5MSA | Gut content | Adult | Sonication | Mobio |
| 5MSL | Gut content | Larvae | Sonication | Mobio |
| 5MSU | Gut content | Substrate | No sonication | Mobio |

\*The number at the beginning of the sample name represents the beetle family group.
The letter after the number is for the method as in column 4: M= Mobio, B= MP, E=
Epicentre. After method: A= Adult, L= larvae, SU= substrate, epA= epithelium adult,
epL= epithelium larvae, SA= sonicated adult, SL= sonicated larvae.

**Supplementary Table 2.**
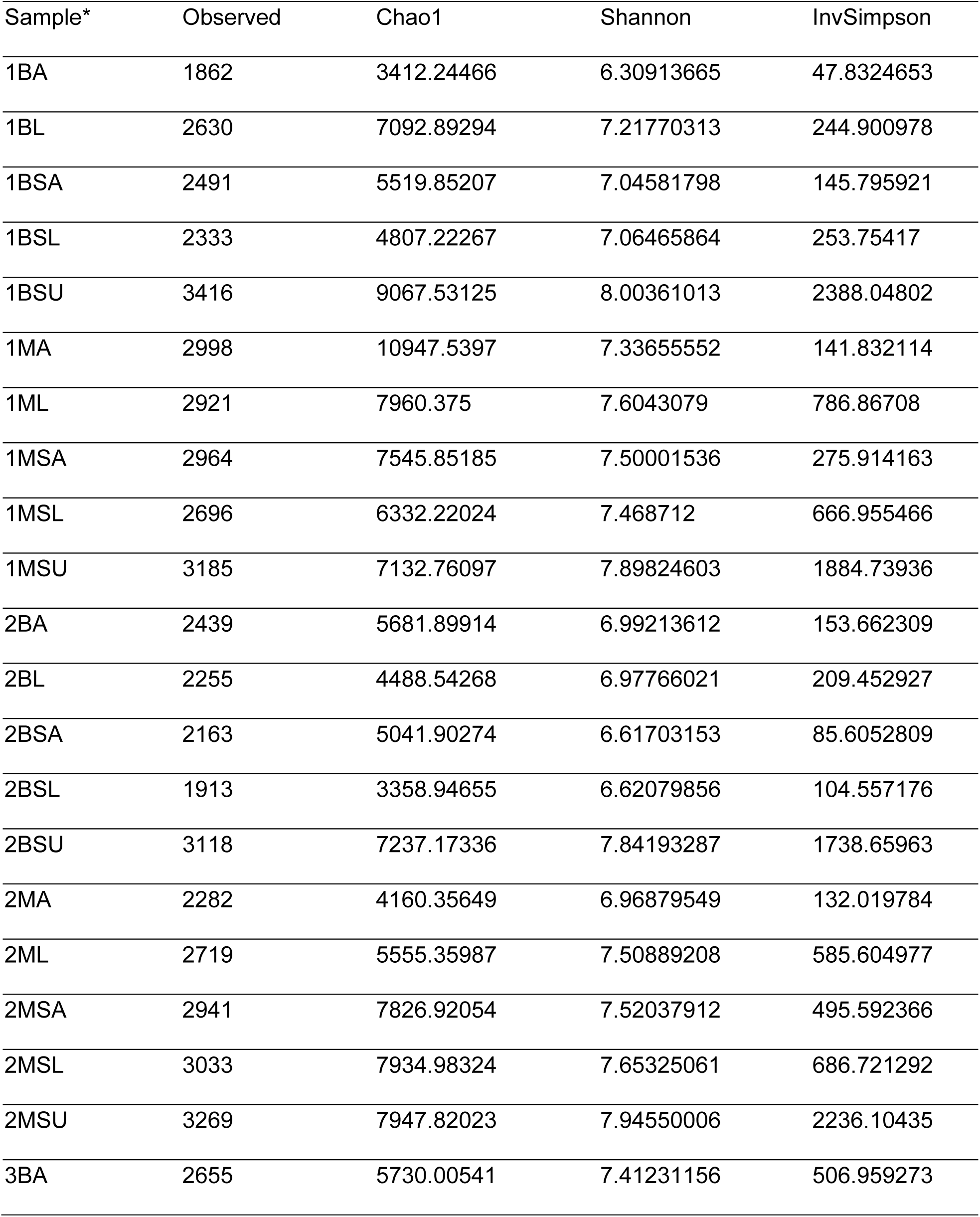
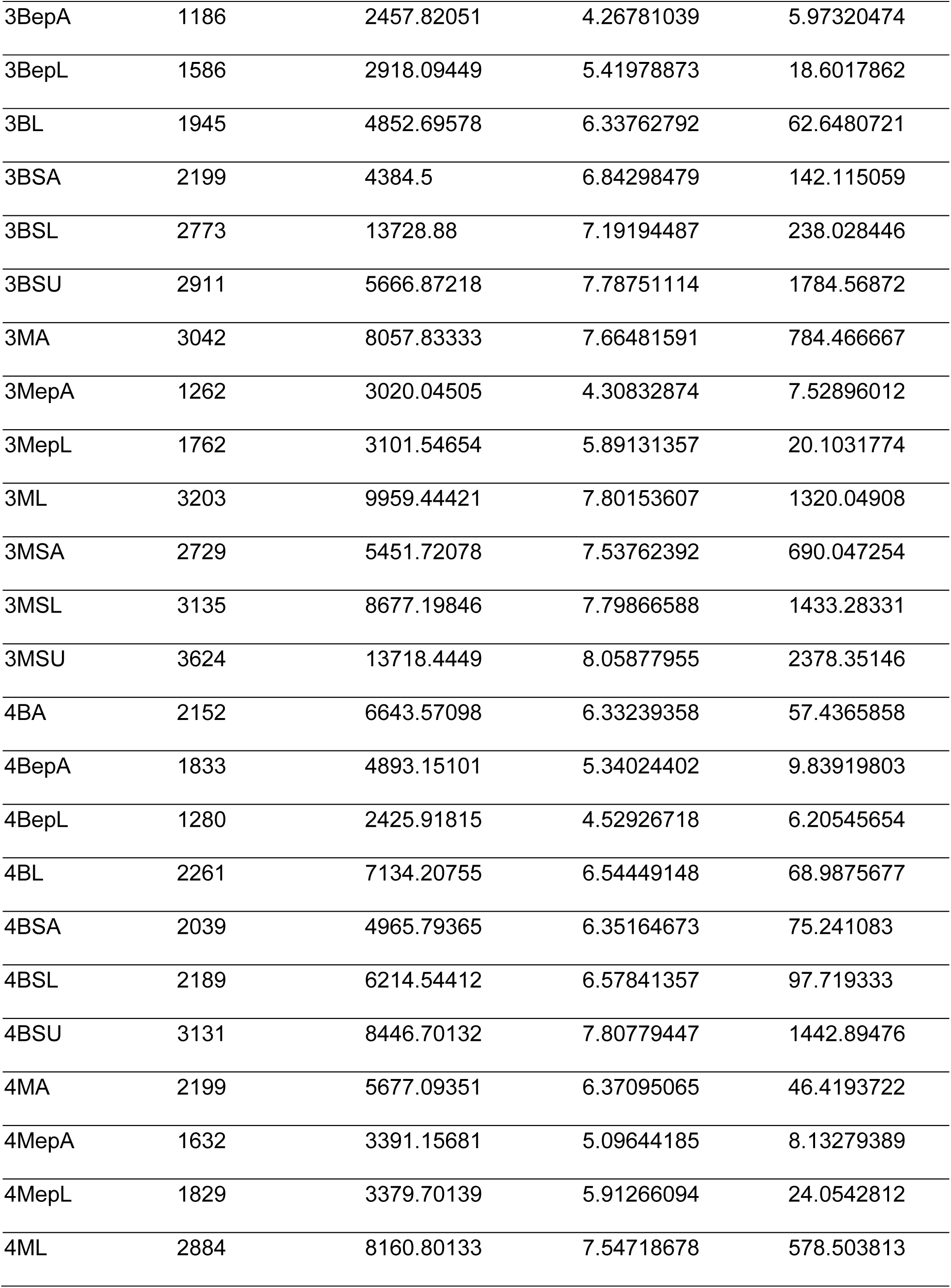
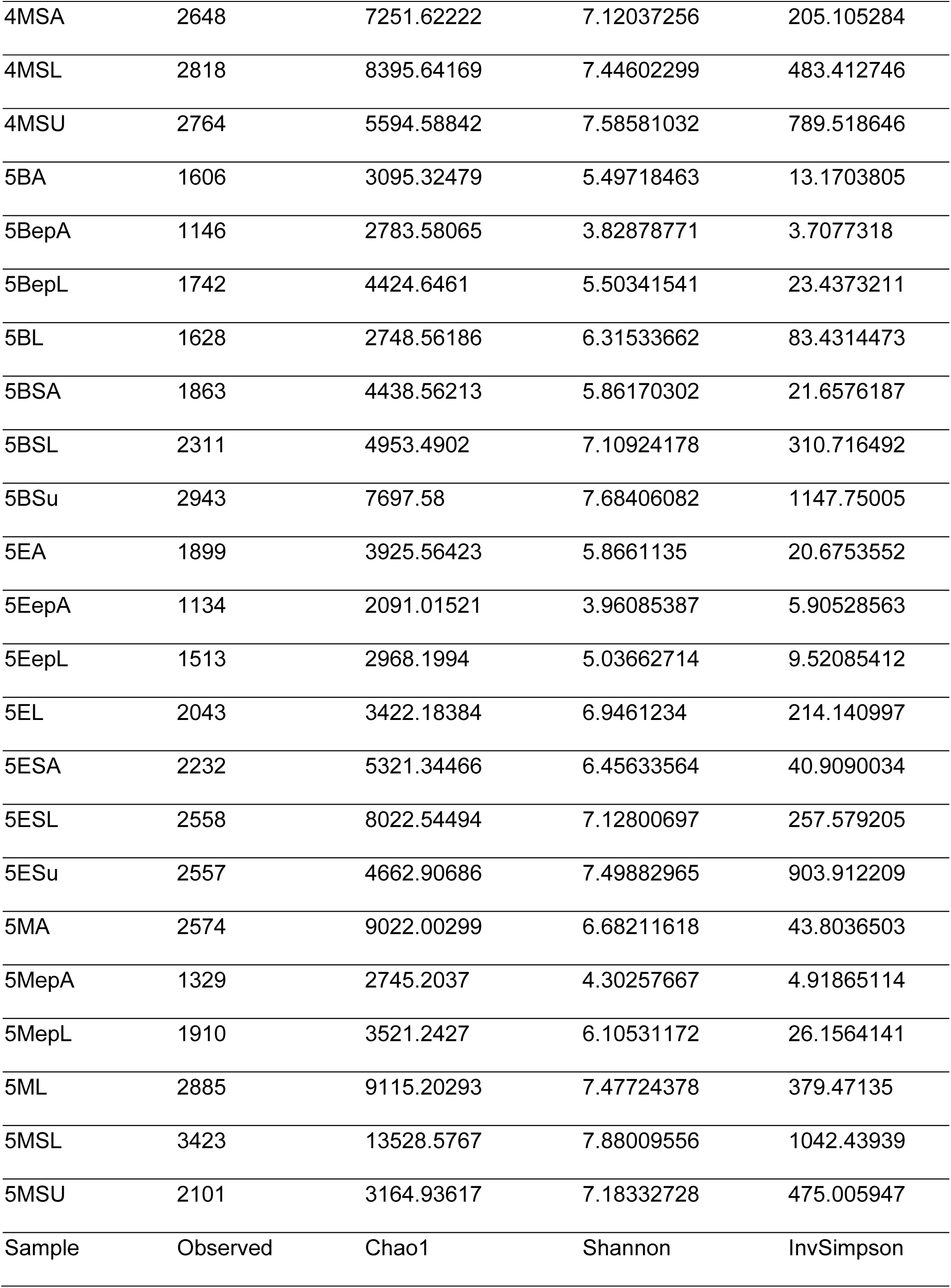
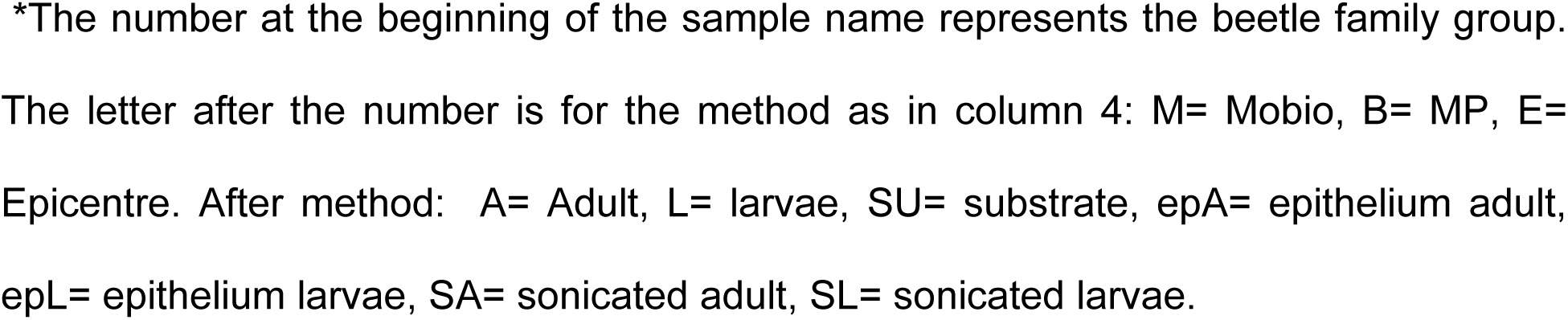
Alpha diversity of the 69 samples of 16S rRNA amplicons from adults, larvae and substrate of the *Veturius* sp.

| Sample* | Observed | Chao1 | Shannon | InvSimpson |
| --- | --- | --- | --- | --- |
| 1BA | 1862 | 3412.24466 | 6.30913665 | 47.8324653 |
| 1BL | 2630 | 7092.89294 | 7.21770313 | 244.900978 |
| 1BSA | 2491 | 5519.85207 | 7.04581798 | 145.795921 |
| 1BSL | 2333 | 4807.22267 | 7.06465864 | 253.75417 |
| 1BSU | 3416 | 9067.53125 | 8.00361013 | 2388.04802 |
| 1MA | 2998 | 10947.5397 | 7.33655552 | 141.832114 |
| 1ML | 2921 | 7960.375 | 7.6043079 | 786.86708 |
| 1MSA | 2964 | 7545.85185 | 7.50001536 | 275.914163 |
| 1MSL | 2696 | 6332.22024 | 7.468712 | 666.955466 |
| 1MSU | 3185 | 7132.76097 | 7.89824603 | 1884.73936 |
| 2BA | 2439 | 5681.89914 | 6.99213612 | 153.662309 |
| 2BL | 2255 | 4488.54268 | 6.97766021 | 209.452927 |
| 2BSA | 2163 | 5041.90274 | 6.61703153 | 85.6052809 |
| 2BSL | 1913 | 3358.94655 | 6.62079856 | 104.557176 |
| 2BSU | 3118 | 7237.17336 | 7.84193287 | 1738.65963 |
| 2MA | 2282 | 4160.35649 | 6.96879549 | 132.019784 |
| 2ML | 2719 | 5555.35987 | 7.50889208 | 585.604977 |
| 2MSA | 2941 | 7826.92054 | 7.52037912 | 495.592366 |
| 2MSL | 3033 | 7934.98324 | 7.65325061 | 686.721292 |
| 2MSU | 3269 | 7947.82023 | 7.94550006 | 2236.10435 |
| 3BA | 2655 | 5730.00541 | 7.41231156 | 506.959273 |
| 3BepA | 1186 | 2457.82051 | 4.26781039 | 5.97320474 |
| 3BepL | 1586 | 2918.09449 | 5.41978873 | 18.6017862 |
| 3BL | 1945 | 4852.69578 | 6.33762792 | 62.6480721 |
| 3BSA | 2199 | 4384.5 | 6.84298479 | 142.115059 |
| 3BSL | 2773 | 13728.88 | 7.19194487 | 238.028446 |
| 3BSU | 2911 | 5666.87218 | 7.78751114 | 1784.56872 |
| 3MA | 3042 | 8057.83333 | 7.66481591 | 784.466667 |
| 3MepA | 1262 | 3020.04505 | 4.30832874 | 7.52896012 |
| 3MepL | 1762 | 3101.54654 | 5.89131357 | 20.1031774 |
| 3ML | 3203 | 9959.44421 | 7.80153607 | 1320.04908 |
| 3MSA | 2729 | 5451.72078 | 7.53762392 | 690.047254 |
| 3MSL | 3135 | 8677.19846 | 7.79866588 | 1433.28331 |
| 3MSU | 3624 | 13718.4449 | 8.05877955 | 2378.35146 |
| 4BA | 2152 | 6643.57098 | 6.33239358 | 57.4365858 |
| 4BepA | 1833 | 4893.15101 | 5.34024402 | 9.83919803 |
| 4BepL | 1280 | 2425.91815 | 4.52926718 | 6.20545654 |
| 4BL | 2261 | 7134.20755 | 6.54449148 | 68.9875677 |
| 4BSA | 2039 | 4965.79365 | 6.35164673 | 75.241083 |
| 4BSL | 2189 | 6214.54412 | 6.57841357 | 97.719333 |
| 4BSU | 3131 | 8446.70132 | 7.80779447 | 1442.89476 |
| 4MA | 2199 | 5677.09351 | 6.37095065 | 46.4193722 |
| 4MepA | 1632 | 3391.15681 | 5.09644185 | 8.13279389 |
| 4MepL | 1829 | 3379.70139 | 5.91266094 | 24.0542812 |
| 4ML | 2884 | 8160.80133 | 7.54718678 | 578.503813 |
| 4MSA | 2648 | 7251.62222 | 7.12037256 | 205.105284 |
| 4MSL | 2818 | 8395.64169 | 7.44602299 | 483.412746 |
| 4MSU | 2764 | 5594.58842 | 7.58581032 | 789.518646 |
| 5BA | 1606 | 3095.32479 | 5.49718463 | 13.1703805 |
| 5BepA | 1146 | 2783.58065 | 3.82878771 | 3.7077318 |
| 5BepL | 1742 | 4424.6461 | 5.50341541 | 23.4373211 |
| 5BL | 1628 | 2748.56186 | 6.31533662 | 83.4314473 |
| 5BSA | 1863 | 4438.56213 | 5.86170302 | 21.6576187 |
| 5BSL | 2311 | 4953.4902 | 7.10924178 | 310.716492 |
| 5BSu | 2943 | 7697.58 | 7.68406082 | 1147.75005 |
| 5EA | 1899 | 3925.56423 | 5.8661135 | 20.6753552 |
| 5EepA | 1134 | 2091.01521 | 3.96085387 | 5.90528563 |
| 5EepL | 1513 | 2968.1994 | 5.03662714 | 9.52085412 |
| 5EL | 2043 | 3422.18384 | 6.9461234 | 214.140997 |
| 5ESA | 2232 | 5321.34466 | 6.45633564 | 40.9090034 |
| 5ESL | 2558 | 8022.54494 | 7.12800697 | 257.579205 |
| 5ESu | 2557 | 4662.90686 | 7.49882965 | 903.912209 |
| 5MA | 2574 | 9022.00299 | 6.68211618 | 43.8036503 |
| 5MepA | 1329 | 2745.2037 | 4.30257667 | 4.91865114 |
| 5MepL | 1910 | 3521.2427 | 6.10531172 | 26.1564141 |
| 5ML | 2885 | 9115.20293 | 7.47724378 | 379.47135 |
| 5MSL | 3423 | 13528.5767 | 7.88009556 | 1042.43939 |
| 5MSU | 2101 | 3164.93617 | 7.18332728 | 475.005947 |
| Sample | Observed | Chao1 | Shannon | InvSimpson |

\*The number at the beginning of the sample name represents the beetle family group. The letter after the number is for the method as in column 4: M= Mobio, B= MP, E= Epicentre. After method: A= Adult, L= larvae, SU= substrate, epA= epithelium adult, epL= epithelium larvae, SA= sonicated adult, SL= sonicated larvae.

**Supplementary Table 3.**
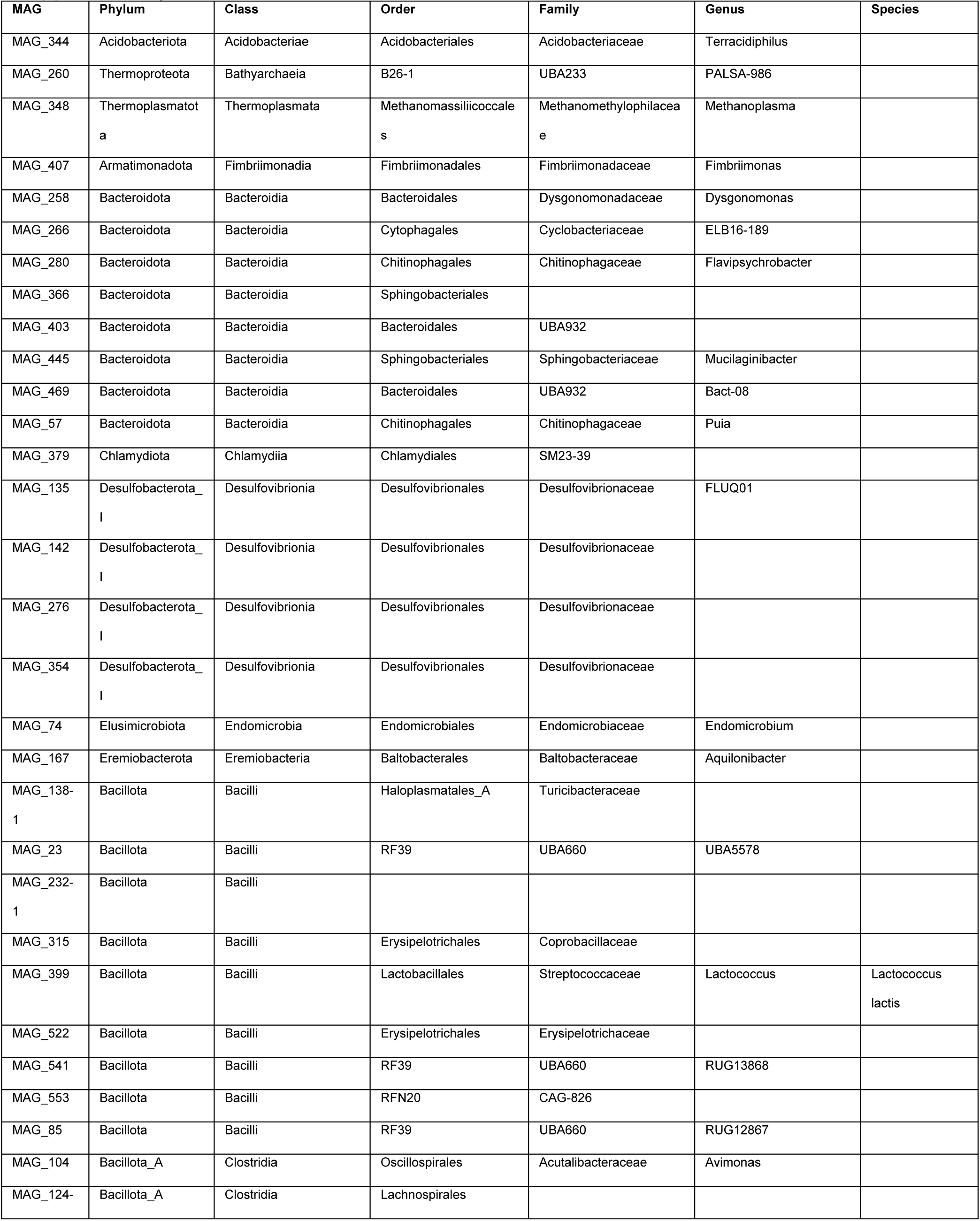
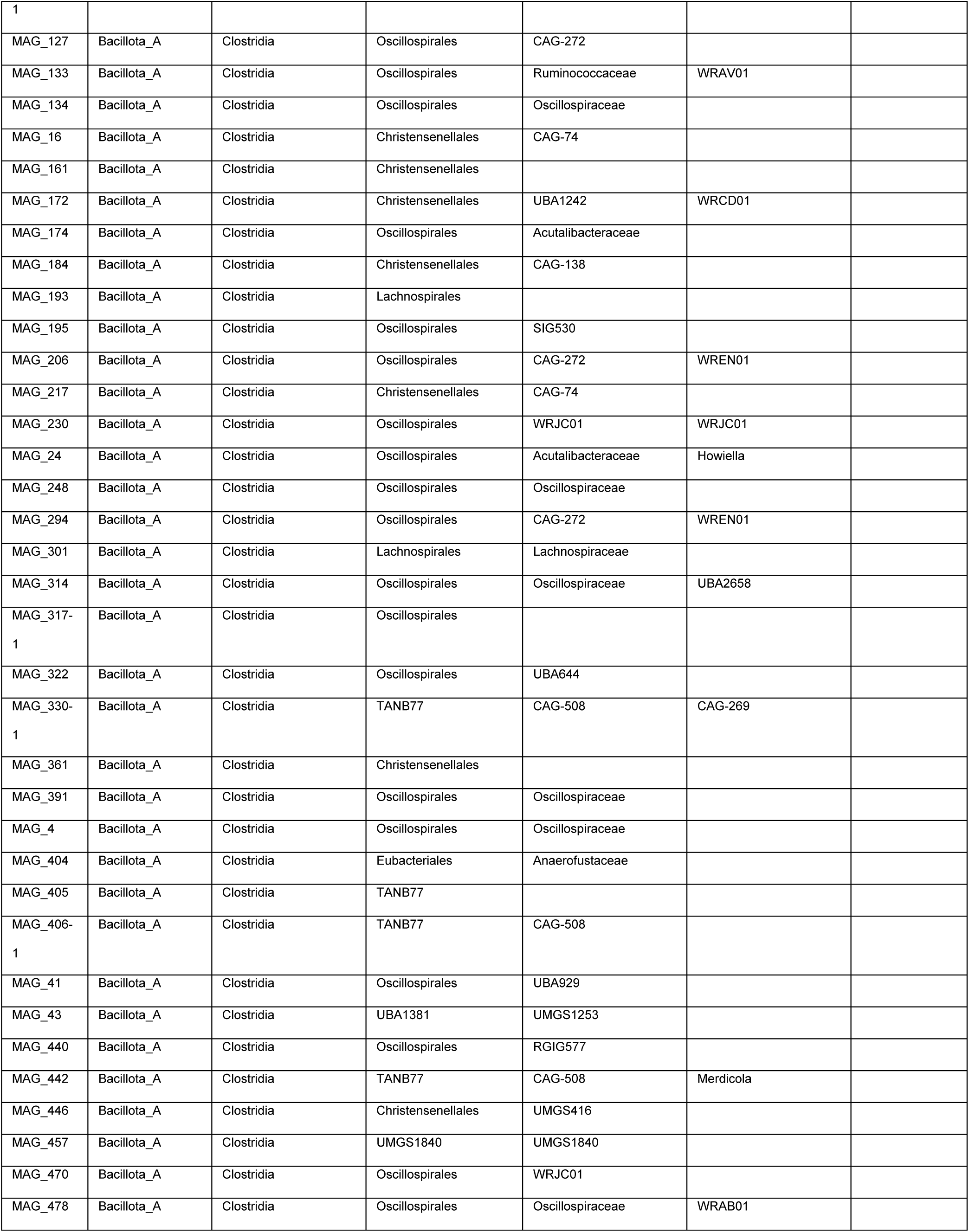
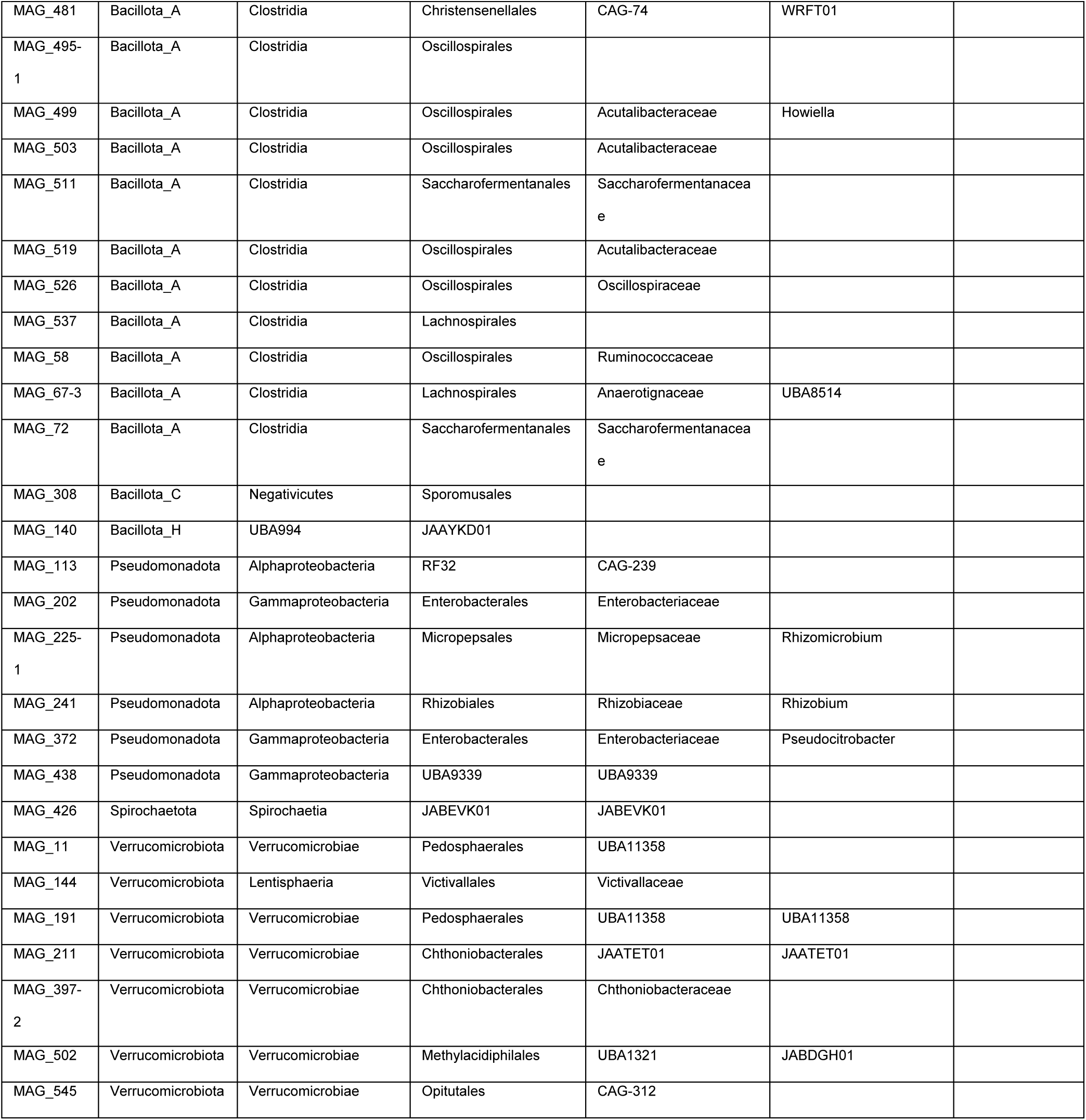
Taxonomic classification of the MAGs.

**Supplementary Figure 1.**
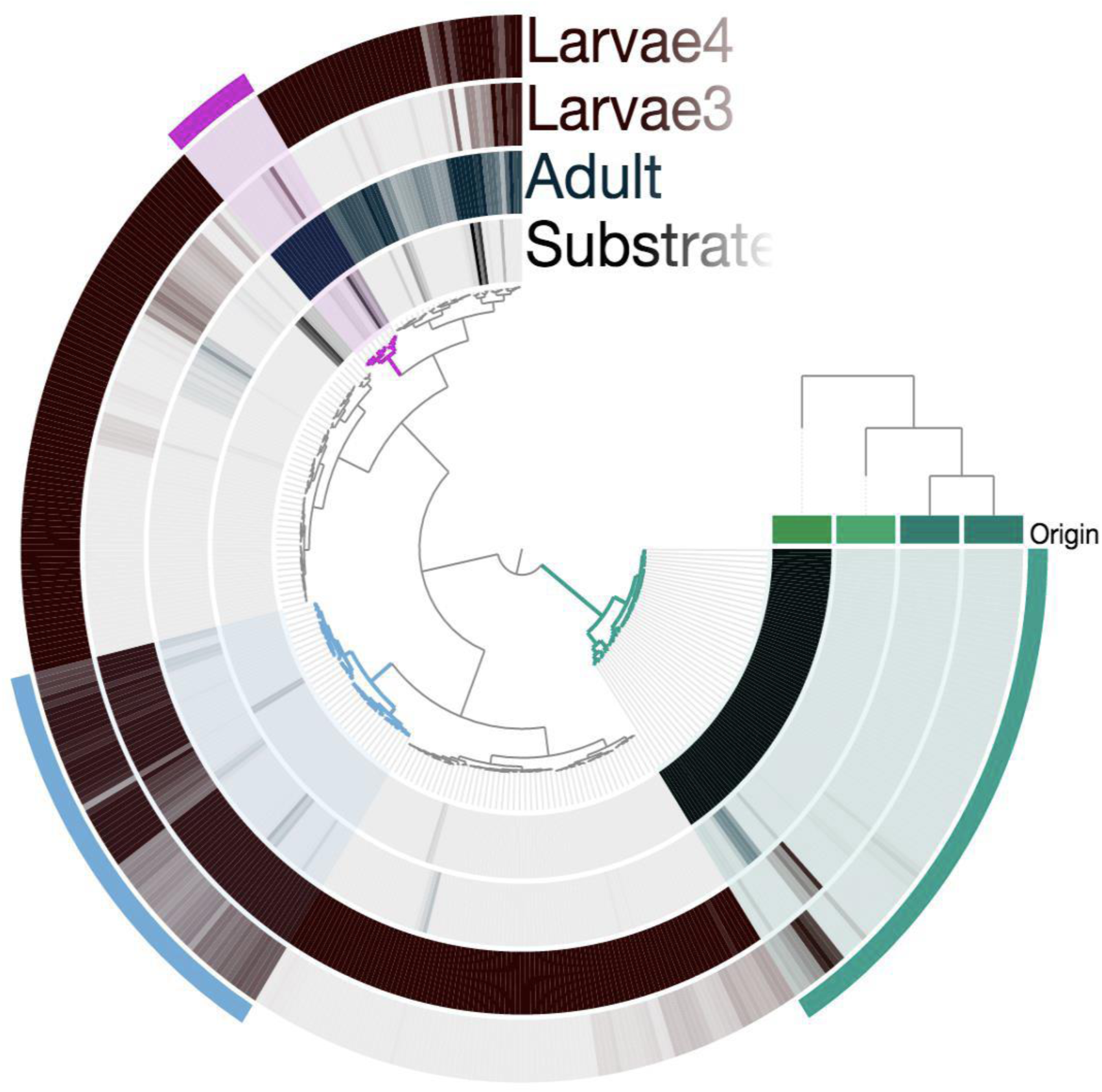
Mapping of 766 bins. Pink branch: metagenomic bins dominant in adult gut. Cyan branch: metagenomic bins enriched in substrate. Blue branch: metagenomic bins found in both larvae samples. Non-colored branch: metagenomic bins dominant in only one larvae sample. The intensity of black of each bin was calculated using Maximum normalized ratios being grey= absent, and black= 100% of bin present in the sample.

**Supplementary Figure 2.**
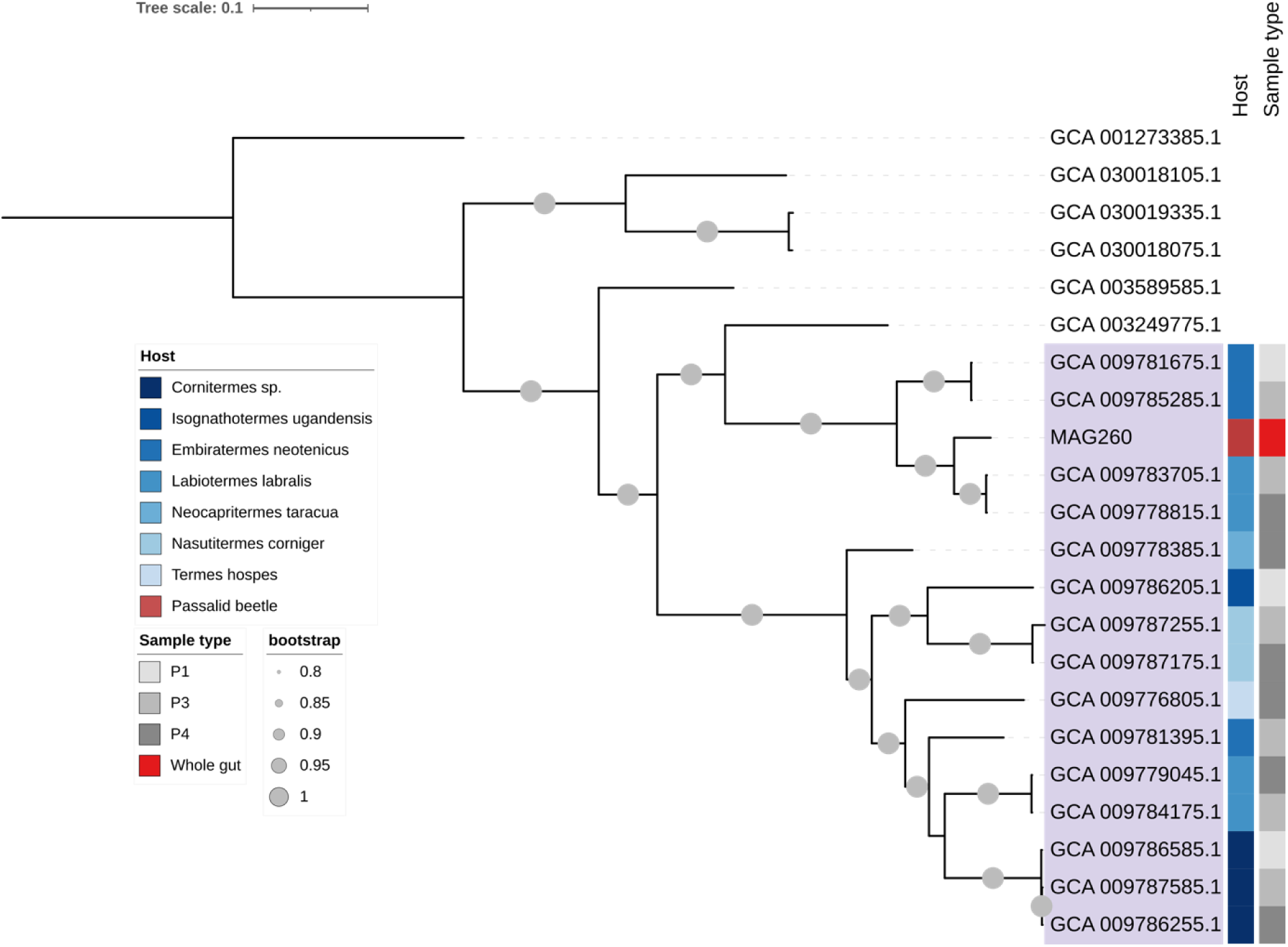
Phylogenomic analysis of the archaeal MAG260 based on 76 single copy core genes. Insect associated genomes are highlighted in purple. The host taxonomy of the insect associated genomes is shown in shades of blue. The Passalid beetle MAG is shown in red. Sample type= segment of the gut from which the genome was reconstructed. Bootstrap values are shown in gray circles, only values >0.8 are shown. Termite Bathyarchaeia genomes were previously published by Loh *et al* 2021^42^.

